# The Role of IgG Subclass in Antibody-Mediated Protection against Carbapenem-Resistant *Klebsiella pneumoniae*

**DOI:** 10.1101/2020.07.24.220780

**Authors:** Michael P. Motley, Elizabet Diago-Navarro, Kasturi Banerjee, Sean Inzerillo, Bettina C. Fries

## Abstract

Monoclonal antibodies (Abs) have the potential to assist in the battle against multidrug-resistant bacteria such as Carbapenem-Resistant *Klebsiella pneumoniae* (CR-*Kp*). However, the characteristics by which these Abs function, such as the role of antibody subclass, must be determined before such modalities can be carried from the bench to the bedside. We performed a subclass switch on anti-capsular monoclonal murine IgG_3_ (mIgG_3_) hybridomas and identified and purified a murine IgG_1_ (mIgG_1_) hybridoma line through sib selection. We then compared the ability of the mIgG_1_ and mIgG_3_ antibodies to control CR-*Kp* ST258 infection both *in vitro* and *in vivo*. We found by ELISA and flow cytometry that mIgG_3_ has superior binding to CR-*Kp* CPS and superior agglutinating ability compared to mIgG_1_. The mIgG_3_ also predictably had better complement-mediated serum bactericidal activity than the mIgG_1_ and also promoted neutrophil-mediated killing at concentrations lower than the mIgG_1_. In contrast, the mIgG_1_ had marginally better activity in improving macrophage-mediated phagocytosis. Comparing their activities in a pulmonary infection model with wild type as well as neutropenic mice, both antibodies reduced organ burden in a non-lethal challenge, regardless of neutrophil status, with mIgG_1_ having the highest overall burden reduction in both scenarios. However, at a lethal inoculum, both antibodies showed reduced efficacy in neutropenic mice, with mIgG_3_ retaining the most activity. These findings suggest the viability of monoclonal Ab adjunctive therapy in neutropenic patients that cannot mount their own immune response, while also providing some insight into the relative contributions of immune mediators in CR-*Kp* protection.

**Importance:** Carbapenem-resistant *Klebsiella pneumoniae* is an urgent public health threat that causes life-threatening infections in immunocompromised hosts. Its resistance to nearly all antibiotics necessitates novel strategies to treat it, including the use of monoclonal antibodies. Monoclonal antibodies are emerging as important adjuncts to traditional pharmaceuticals, and studying how they protect against specific bacteria such as *Klebsiella pneumoniae* is crucial to their development as effective therapies. Antibody subclass is often overlooked but is a major factor in how an antibody interacts with other mediators of immunity. This paper is the first to examine how the subclass of anti-capsular monoclonal antibodies can affect efficacy against CR-*Kp*. Additionally, this work sheds light on the viability of monoclonal antibody therapy in neutropenic patients, who are most vulnerable to CR-*Kp* infection.

## INTRODUCTION

Monoclonal antibodies (Abs) are becoming increasingly important in the treatment of a variety of different diseases, including infectious disease (1, 2). The escalating failure of traditional antibiotics to treat bacterial infections further emphasizes the importance of testing alternative therapies, including monoclonal Abs, against these pathogens (3). Much information regarding how Ab structure influences interactions with pathogens remains to be discovered, and until recently, the role of the Ab constant region and its different variants, or subclasses, had often been overlooked in therapeutic monoclonal Ab development. While four subclasses of IgG Abs exist in humans, the majority of monoclonal Abs used in the clinic are human IgG1 (hIgG_1_), the most prevalent subclass (4). The subclass of an Ab, dictated by the number of disulfide bonds joining the heavy chains; its fragment crystallizable (Fc) region; and other aspects of the heavy chain, affects what immune receptors and adaptors the antibody binds. Subsequently, these interactions determine the amplitude and character of the immune response (5). Some subclasses interact with more immunostimulatory Fc receptors on professional phagocytes, increasing their activity, while others bind to immunosuppressing receptors that act to reduce collateral damage caused by excessive inflammation (6). Additionally, subclasses can be responsible for differences in antigen binding, even when Abs have identical variable regions (7, 8). Understanding differences between IgG subclasses – how they bind, interact with the pathogen, and interact with other facets of immunity - is important to both understanding which subclasses may provide a therapeutic benefit (8–12).

With the recent rise of multidrug resistant gram-negative bacteria, such as carbapenem-resistant *Klebsiella pneumoniae* (CR-*Kp*) (13), several laboratories have been focusing on developing antibodies against these pathogens (3, 14–17). These bacteria frequently infect immunocompromised populations that lack robust innate and adaptive immune responses (18, 19). Therefore, it is crucial to understand not only how different Ab subclasses act against these pathogens, but also how they function in the context of immunocompromised states. We recently cloned murine IgG_3_ (mIgG_3_) monoclonal Abs that targeted the capsular polysaccharide (CPS) of *wzi154* CR-*Kp* isolates, which fall within the clade 2 subfamily of the CR-*Kp* ST258 clonal group (14). Isolates of this conserved subgroup have been shown to be susceptible to Ab therapy through a variety of *in vitro* modalities such as killing by serum complement and action by neutrophils and macrophages, and such antibodies have been shown to be protective *in vivo* as well (14, 15, 20). We chose one of these, 17H12, to study the effects of switching IgG subclass on anti-*Klebsiella* Ab functionality. We report findings that the parent mIgG_3_ was superior to the new murine IgG_1_ (mIgG_1_) variant in binding ability, initiation of complement-mediated bactericidal activity by serum, and activation of neutrophil-mediated killing at lower antibody concentrations. Conversely, the new mIgG_1_ variant slightly outperformed the mIgG_3_ parent in promoting macrophage-mediated phagocytosis of the bacteria. Finally, our comparison within a pulmonary mouse challenge model shows comparable overall efficacy of both subclasses in reducing bacterial organ burden at both a lethal and non-lethal inoculum in wild type mice. Interestingly efficacy of both antibodies was maintained in neutropenic mice except at the lethal inoculum.

## RESULTS

### Parent 17H12 mIgG_3_ variant has superior binding of *wzi154* capsular polysaccharide relative to new subclass switch variant 17H12 mIgG_1_ despite identical variable regions

We first isolated a mIgG_1_ variant of the mIgG_3_ 17H12 hybridoma line by using sib selection followed by FACS and soft agar cloning, which we previously utilized (12). Though we sought to generate all three additional subclasses, only mIgG_1_ and mIgG_2a_ variants were discovered in our initial screen, and only mIgG_1_ variants could be enriched by downstream sib selection. The mIgG_1_ hybridomas were verified to exclusively produce mIgG_1_, and the sequence of the variable region of the new clone was found to be identical to that of the mIgG_3_ parent (the characteristics of which have been previously published (14)).

To investigate how subclass switching affected binding, we compared the affinity of the new mIgG_1_ relative to its parent mIgG_3_ against the *wzi154* CPS originally used to generate the mAb (14). Analysis by ELISA showed the mIgG_1_ to have 4-fold less binding than its mIgG_3_ counterpart, with EC_50_ values of 27.6nM (95% CI 18.8nM-40.6nM) and 6.81nM (95% CI 3.00nM-13.2nM), respectively (**Figure 1A**).

**Figure 1.**
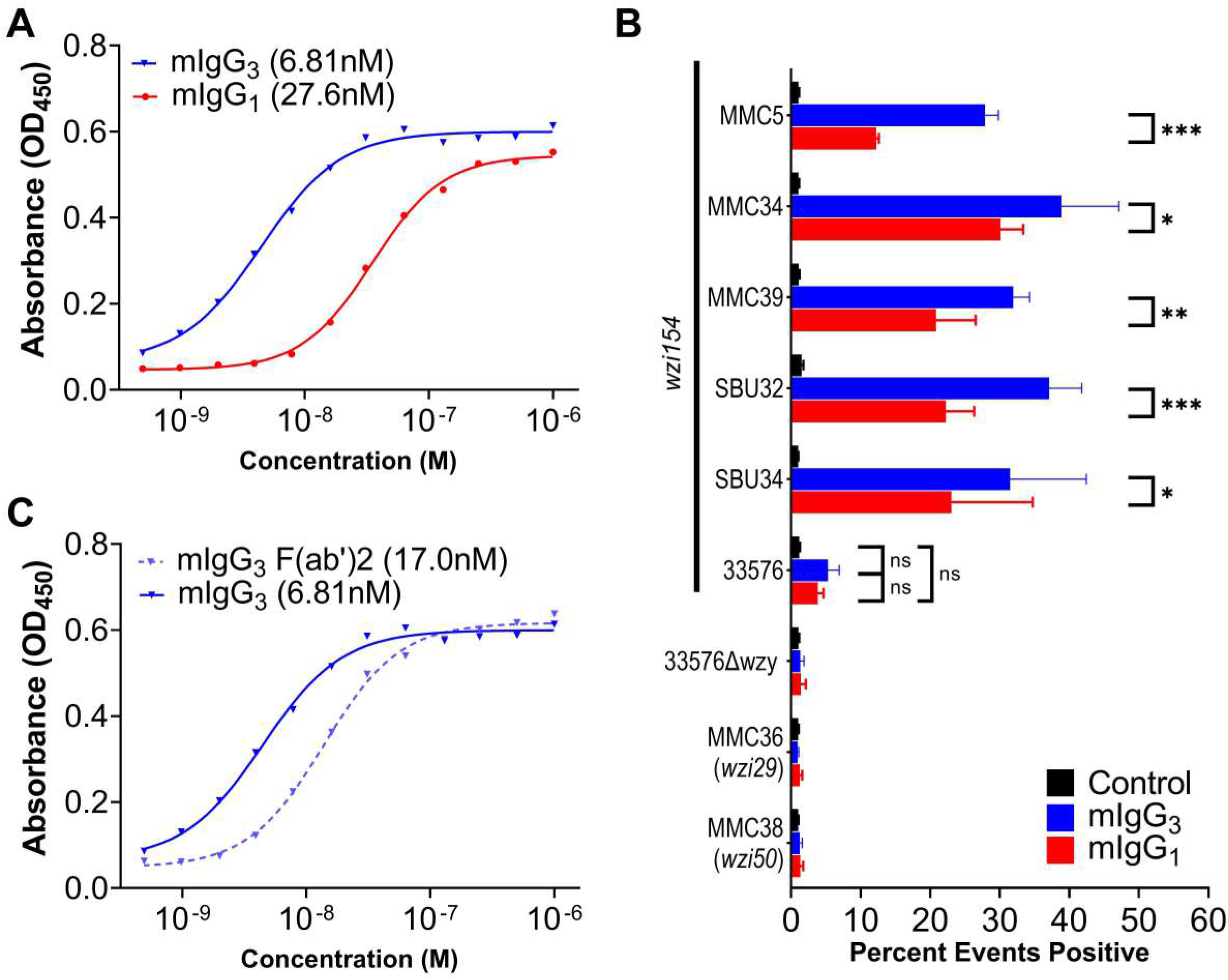
Comparison of binding and agglutination of 17H12 mIgG_1_ and mIgG_3_. (A) Binding curves of 17H12 mIgG_1_ and m17H12 IgG_3_ measured by indirect ELISA on plates coated with *wzi154* (MMC34) CPS. EC_50_ values are displayed with legend. The plot is representative of four independent experiments. Differences by unpaired t test were determined to be significant (p=0.0013) (B) Relative agglutination of nine CR-*Kp* strains measured by flow cytometry. Agglutination was measured as percentage of events that exceeded the maximum forward scatter of bacteria without antibody (Percent positive). Error bars indicate the SD from three independent experiments. Two-Way ANOVA determined a significant difference between treatment groups across strain types (p<0.001), with results of individual comparisons using Tukey’s post-hoc test displayed in-graph: p< 0.05 (*),<0.01 (**), p< 0.001 (***). (C) Binding curves of 17H12 mIgG_3_ and its F(ab’)_2_ fragment, measured by indirect ELISA. The plot is representative of four independent experiments. EC_50_ values are displayed with legend. Differences between EC_50_ values were determined to be significant by unpaired t test (p<0.01).

Next, we compared the ability of each Ab to agglutinate CR-*Kp* clinical isolates, utilizing flow cytometry to measure relative clump sizes by forward scatter (21). We began testing the Abs with the previously-studied CR-*Kp wzi154* strain 39 (MMC39) (14) which we transformed with a novel GFP-expressing plasmid pProbe-KtBl. Using this transformant, referred to hereafter as MMC39-GFP, we noted that mIgG_3_ promoted better agglutination than mIgG_1_; mIgG_3_-opsonized bacteria demonstrated higher forward scatter than mIgG_1_-opsonised bacteria at the same concentration of antibody and achieved maximum forward scatter at lower concentrations of antibody as well. (**Figure S1**). As 30μg/mL provided the greatest disparity in agglutination between mIgG_1_ and mIgG_3_, we chose this antibody concentration and then compared relative agglutination across a number of CR-*Kp* isolates **(Table 1**). These isolates include those collected from Montefiore Medical Center (MMC) and Stony Brook University (SBU), including those previously studied (14, 22), as well as a previously-studied isolate 33576 from the mid-Atlantic states and its capsule-deficient variant (15) (**Table 1**). These strains cover the three most prevalent *wzi* subgroups within the ST258 clone (22–24). Measuring the percentage of aggregates larger than a baseline formed by control bacteria untreated with antibody (25), we found that both Abs improved agglutination of nearly all *wzi154* strains relative to a control mIgG_1_, but did not significantly promote agglutination of capsule-deficient strain 33576Δwzy, or CR-*Kp* isolates of the *wzi29* and *wzi50* capsule types (**Figure 1B**). Additionally, at this concentration the mIgG_3_ parent caused higher percent-above-baseline agglutination than mIgG_1_ in 5 of 6 isolates. While percent agglutination of strain 33576 did not significantly improve above baseline for either 17H12 mIgG_1_ or mIgG_3_, this was likely due to high observed baseline aggregation of the strain. In contrast, histograms of gated 33576 (and MMC5) demonstrate clear shifts in forward scatter between the mIgG_3_, mIgG_1_, and control groups, and aggregated raw mean forward scatter data show significantly higher agglutination by either Ab of all *wzi154* bacteria relative to both controls (**Figure S1**).

**Table 1:** List of CR-*Kp* strains used in the study

| Strain | ST | *wzi* | Source of Isolate | Ref |
| --- | --- | --- | --- | --- |
| MMC5 | 258 | 154 | Blood isolate from MMC | (14) |
| MMC39 | 258 | 154 | Blood isolate from MMC | (14) |
| MMC39-GFP | 258 | 154 | Transformant of MMC39 |  |
| SBU32 | 258 | 154 | Urine isolate from SBU | (14) |
| SBU34 | 258 | 154 | Urine isolate from SBU | (14) |
| 33576 | 258 | 154 | Isolate from Mid Atlantic | (20) |
| 33576Δwzy | 258 | n/a | Isogenic Mutant of 33576 | (15) |
| MMC36 | 258 | 29 | Blood isolate from MMC | (14) |
| MMC38 | 258 | 50 | Blood isolate from MMC | (14) |

As previous studies have shown the importance of the Fc region in Ab binding to antigen, we performed F(ab’)_2_ digests of both Abs to determine whether this may hold true for 17H12. We determined by ELISA that digestion of 17H12 mIgG_3_ led to a 2.7-fold loss of binding relative to whole mIgG_3_ (**Figure 1C**). In contrast, digestion of 17H12 mIgG_1_ did not impact binding (**Figure S2**).

### Complement-dependent serum killing was exclusive to mIgG_3_

We next compared the ability of the two subclasses to mediate serum killing of CR-*Kp*. Using 20% fresh human serum, we determined that parent mIgG_3_ caused a 90.1% and 92.7% reduction in CFU of MMC39 CR-*Kp* after 60 and 120 min, respectively. In contrast, mIgG_1_ caused a 64.4% and 63.5% drop in CFU, similar to the control Ab (**Figure 2A**). In strain 33576 20% human sera was found to be insufficient to reduce CFU under any condition **(Figure 2B**), but bacterial replication was inhibited by both subclasses, while bacteria exposed to the control multiplied 6-fold over 120 min. Increasing the percentage sera to 40% improved CFU reduction of 33576, but in an antibody-independent manner **(Figure S3**). As previously described, 33576Δwzy exhibits pronounced sensitivity to serum in the absence of capsule (15), and near-complete killing occurred irrespective of treatment **(Figure S3)**. Variability in killing was observed depending on the human serum donor, but effects were consistent between experiments using serum from the same donor. Additionally, heat-inactivation (HI) of serum abrogated all killing effects against MMC39, and instead allowed *Kp* growth, which both Abs partially limited **(Figure 2A**).

**Figure 2.**
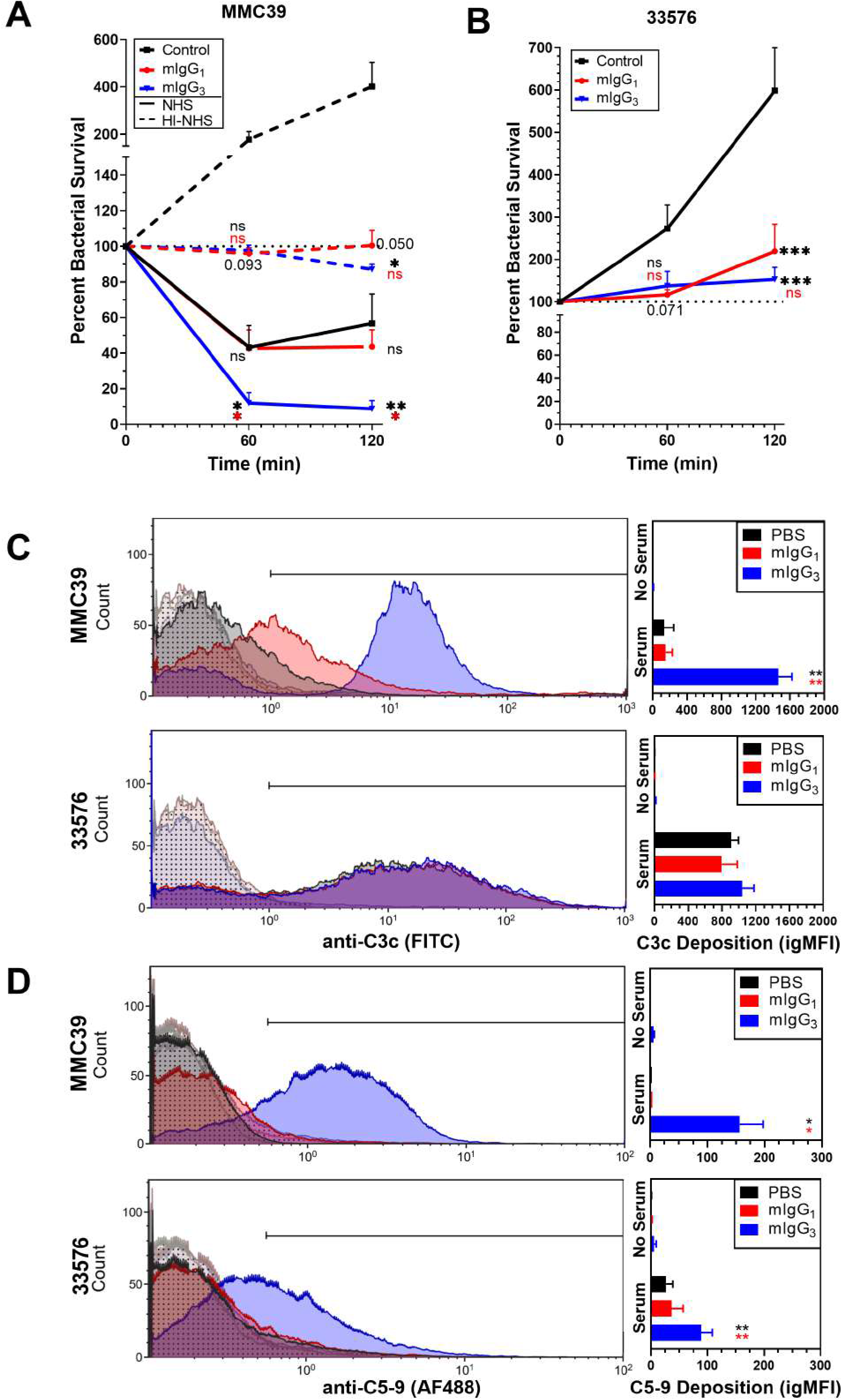
Serum Bactericidal Effect and Complement Deposition mediated by 17H12 mIgG_1_ and mIgG_3_. (A, B) Growth curves of MMC39 (A) and 33576 (B) in 20% NHS (both, solid lines) or in 20% HI-NHS (A only, dashed lines) supplemented with 40μg/mL of indicated mAb. 100% represents no increase in CFU from baseline. Error bars indicate SEM of the at least five independent experiments for MMC39, three for 33576. Overall differences between treatment groups in MMC39 (A) and in 33576 (B) were determined to be significant by Repeated-Measures Two Way ANOVA (p= 0.0380 and 0.0124, respectively) with results of individual comparisons at 60 min and 120 min using Tukey’s post-hoc test displayed in-graph. (C, D) Fixation of complement components onto MMC39, and 33576, as measured by flow cytometry. Left histogram overlays depict representative data from three independent experiments, and right graphs show the means and SEM of the integrated geometric mean fluorescence intensity (igMFI) of each experiment. Histograms of treatments with serum are filled with solid colors, while those without serum have patterned fills. Overall differences between the variances of all treatments for each strain and each complement component were assessed for significance by Repeated Measures One-Way ANOVA (MMC39 C3c, p=0.002, 33576 C3c p=0.228, MMC39 C5-9 p=0.013, 33576 C5-9 p=0.020) with results of Tukey’s post-hoc test for multiple comparisons displayed in-graph. For all in-graph statistics, p values displayed in black are comparisons to the control IgG, whereas p values in red compare mIgG_1_ with mIgG_3_. p values are replaced with ns if >0.1 (not significant); * if p < 0.05; ** if p < 0.01; and *** if p <-0.001.

We also specifically compared the relative amount of complement each antibody could fix. Using flow cytometry we detected C3c and C5b-9 Membrane Attack Complex deposition on MMC39, 33576, and 33576Δwzy in the presence of either subclass **(Figure 2C,D, Figure S3)**. We observed the parent mIgG_3_ to outperform the PBS control and the mIgG_1_ in deposition of C3c onto MMC39, but not onto 33576, which exhibited high background C3c deposition **(Figure 2C)**. With both strains however, C5b-9 deposition was found to be increased when bacteria were pre-opsonized with mIgG_3_ **(Figure 2D)**. Controls confirmed that mIgG_3_ capsule binding caused deposition of serum-based complement, with incubation of the bacteria in 0% (**Figure 2C,D)** or 20% HI Serum (**Figure S3**) resulting in no detectable deposition, and the antibody failing to deposit additional complement onto the capsule-deficient 33576Δwzy **(Figure S3)**.

### Both subclasses improved macrophage mediated phagocytosis, with mIgG_1_ performing marginally better than mIgG_3_

We next compared the ability of the subclasses to contribute to cell-mediated action against CR-*Kp*. Monocytes and macrophages are important in CR-*Kp* clearance (26), and we have previously demonstrated 17H12 mIgG_3_ to enhance phagocytic uptake of numerous *wzi154* CR-*Kp* strains (14). Therefore, we compared the ability of both variants to promote uptake using a CFU-based phagocytosis assay we previously performed (14, 27). Our data show mIgG_1_ to slightly improve J774A.1 macrophage phagocytosis of MMC39 relative to mIgG_3_, a trend that was also suggested in the phagocytosis of 33576 **(Figure 3A)**. In contrast, 33576Δwzy was phagocytosed irrespective of treatment condition. The difference in phagocytosis between mIgG_1_ and mIgG_3_ was subtle, and not evident when phagocytosis of MMC39-GFP was observed under fluorescent microscopy **(Figure 3B)**. Additionally, we found that uptake was not correlated with intracellular killing; after both J774A.1 macrophages and bone marrow derived macrophages had phagocytized the opsonized bacteria, and external bacteria had been washed away, we observed by both CFU quantitation and by microscopy that the number of bacteria within these cells increased over time (data not shown). This observation suggests intracellular multiplication of CR-*Kp* after phagocytosis.

**Figure 3.**
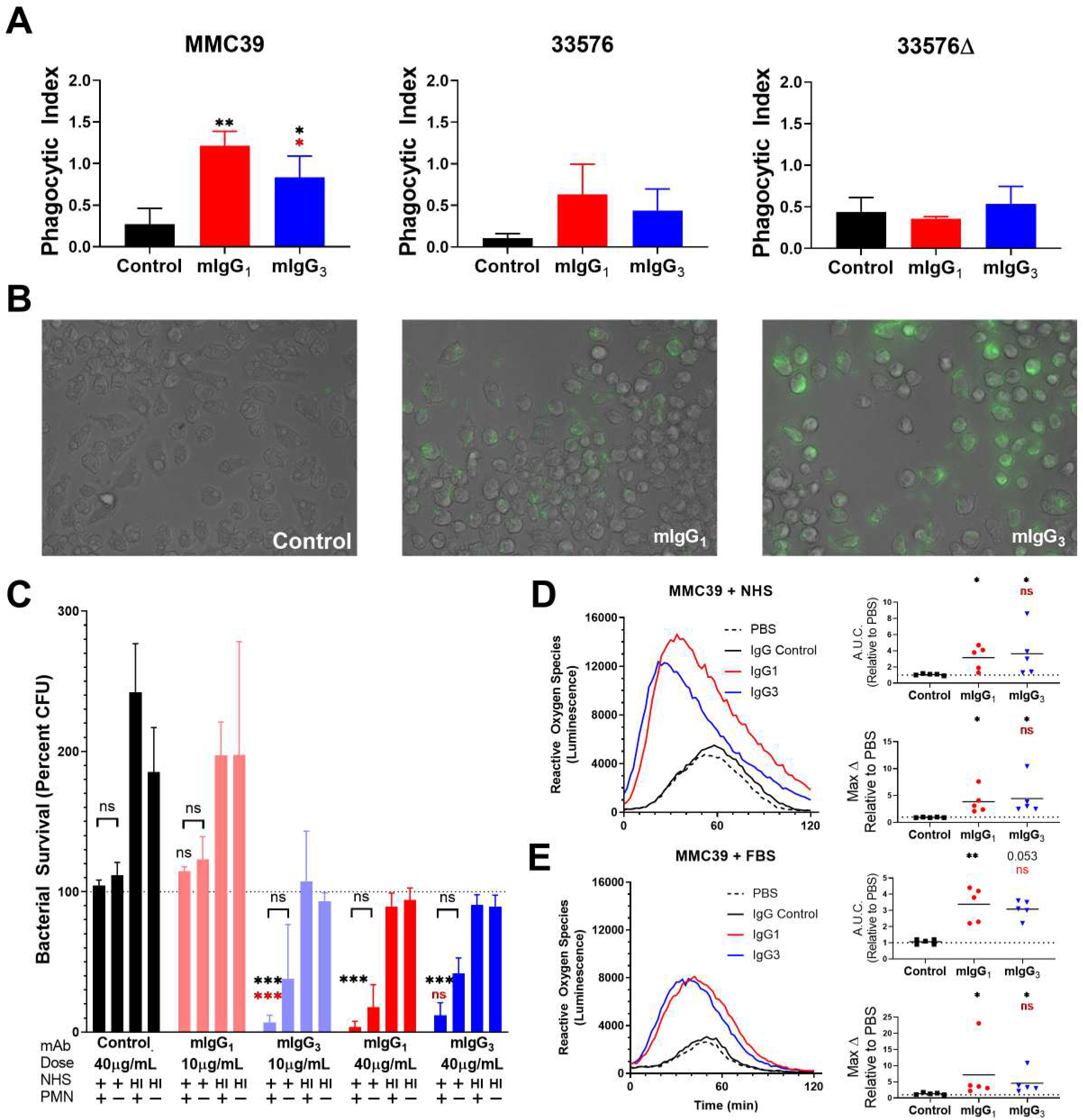
Comparison of cell-mediated phagocytosis and killing. (A) Phagocytosis of MMC39, 33576, and 33576Δ by J774.A1 murine macrophage-like cells after incubation with 40μg/mL of respective antibody. Phagocytic index is calculated as the number of CFU recovered from the plate, divided by the number of cells plated. Bars depict means and SEMs of three independent experiments, with wells performed in triplicate. Overall differences in the variance of all treatments for each strain were assessed for significance by Repeated Measures One-Way ANOVA (39 p<0.001, 33576 p=0.221, 33576Δwzy p=0.520), with results of Tukey’s post-hoc test for multiple comparisons displayed in-graph. (B) Representative images of antibody-mediated phagocytosis of MMC39-GFP by J774A.1 macrophages. Images were taken at 40x with an EVOS microscope using brightfield and GFP channels. (C) Killing of pre-opsonized MMC39 by human neutrophils after 60 min. Bars depict Mean and SEM of three independent experiments. Within the results of the NHS-treated samples, differences in the variance of dose-matched treatment groups with and without neutrophils were assessed for significance by Two-Way Repeated-Measures ANOVA (Variance between treatment groups p<0.001 for both 10μg and 40μg sets; Variance comparing neutrophil status p=0.135 and p=0.058, respectively) with results of Sidak’s multiple comparisons tests displayed in-graph. (D, E) Reactive Oxygen Species production by human neutrophils exposed to pre-opsonized MMC39, as measured by luminol luminescence, in the presence of NHS (D) or FBS (E). The left time lapse graphs graphs are representative of five independent experiments.. Right bar graphs show aggregate data of the Area Under the Curve (A.U.C.) and the maximum rate of change (Max Δ), relative to PBS, for all experiments. Differences in A.U.C. and Max Δbetween control Ab, mIgG_1_ and mIgG_3_ were assessed for significance by a Kruskal-Wallis test (p<0.01 for NHS and FBS), with results of Dunn’s test for multiple comparisons displayed in-graph.For all in-graph statistics, p values displayed in black are comparisons to the control IgG, whereas p values in red compare mIgG_1_ with mIgG_3_. p values are replaced with ns if >0.1 (not significant); * if < 0.05; ** if <0.01; and *** if p< 0.001

### Neutrophil killing of CR-*Kp* improved with lower concentrations of mIgG_3_ than mIgG_1_

We next compared the ability of both antibodies to promote killing by neutrophils. Data in humans has shown that neutropenic patients may have reduced survival in cases of bacteremia caused by CR-*Kp* and other carbapenem-resistant Enterobacteriaceae (28). In mice, some studies have shown neutrophils to be important in CR-*Kp* clearance (15, 29), while others have shown them to be less valuable (26). Using a tube-based incubation assay with human neutrophils, we observed both antibodies to promote neutrophil-dependent killing of MMC39 in the presence of 5% autologous serum at 40μg/mL **(Figure 3C)**. When we reduced the dose to 10μg/mL however, mIgG_3_ demonstrated improved efficacy, while mIgG_1_ lost all efficacy relative to the control. Antibody-mediated killing by neutrophils was dependent on serum, as co-incubation of neutrophils with HI serum failed to reduce bacterial CFU.

We then compared the ability of the two antibodies to promote neutrophil reactive oxygen species (ROS) production in response to CR-*Kp*. We used MMC39, MMC5, SBU32, and SBU34, and either 5% NHS or 20% HI-FBS in reactions (**Figure3D, Figure S4)**. These data indicated that both mIgG_1_ and mIgG_3_ promoted ROS production comparably. However, strains had varying ability to induce ROS production, with strains SBU32 and SBU34 having high constitutive production of ROS in the absence of either monoclonal Ab, whereas base ROS in MMC39 and MMC5 were nearly absent **(Figure3D, Figure 4)**. Furthermore, baseline ROS production in SBU32 and SBU34 occurred irrespective of whether NHS or HI-FBS was used, though NHS appeared to promote marginally higher production.

**Figure 4.**
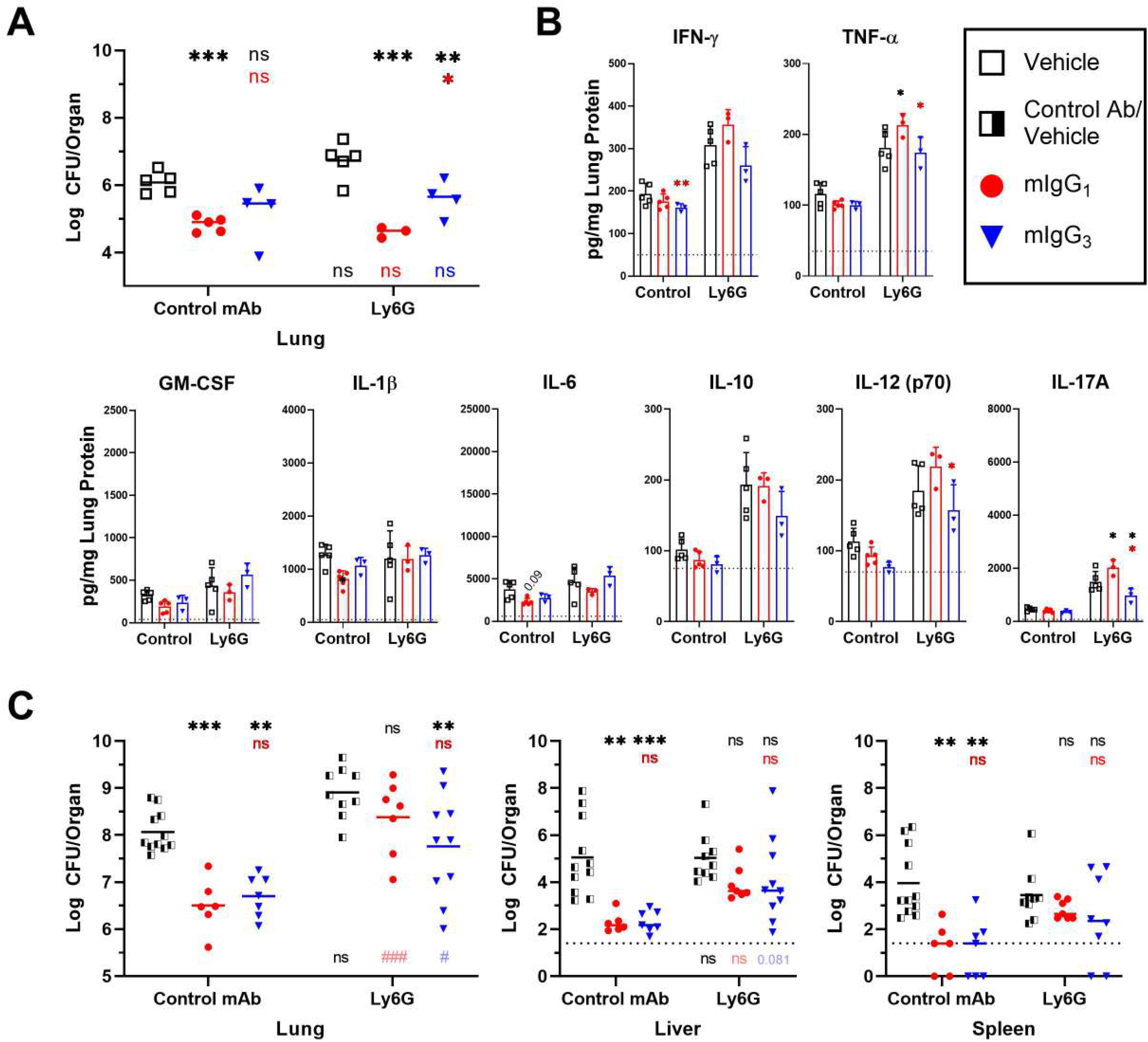
Antibody-mediated protection against CR-*Kp* pulmonary infection in control and neutrophil-depleted mice. (A) Bacterial burden in the lungs of C57Bl/6 depleted of neutrophils (Ly6G) or administered a control antibody and subsequently infected with a non-lethal inoculum of MMC39 pre-opsonized with 17H12 subclasses or controls. Each symbol represents one mouse. (B) Cytokine levels, normalized to total lung protein, of mice in (A), measured by a Bio-Plex panel. (C) Bacterial burden in lungs, liver, and spleen of depleted and non-depleted mice infected with a lethal inoculum of pre-opsonized MMC39. For all studies, overall differences in CFU and cytokines between treatment groups and between neutrophil status were assessed for significance by Two Way-ANOVA. Individual comparisons made between treatment groups of mice of the same neutrophil status (* symbols above), or comparisons made between wild type or neutropenic mice given the same inoculum (# symbols below) were tested using Tukey’s post-hoc test with p values displayed in-graph. p values are replaced with ns if >0.1 (not significant); * if p < 0.05; ** if p < 0.01; and *** if p <-0.001. p values below plots compare CFU or cytokine levels of wild type and neutropenic mice within the same treatment group.

### Both subclasses improved CR-*Kp* lung clearance *in vivo*, including within neutropenic mice

Finally, we compared the protective efficacy of the subclasses *in vivo*, using a pulmonary infection model we previously utilized in BALB/c mice (14). Because we observed these two antibodies to enhance neutrophil-mediated killing, we compared the relative abilities of the subclasses to control organ burden in both c57BL/6 wild type mice as well as neutropenic mice, which were generated by Ly6G antibody-mediated depletion. After depleting neutrophils or administering a control Ab, we infected mice with a sublethal dose of MMC39 pre-opsonized with either mIgG_1_, mIgG_3_, a control mIgG_1_, or Tris-Glycine buffer alone. As we found no significant differences between the mIgG control and the Tris-Glycine vehicle, these groups were combined for analysis. We first observed higher CFU lung burden in neutropenic mice, in contrast to previous studies (26, 30). Additionally, we observed efficacy of both subclasses in reducing bacterial burden in the lung in both wild type and in neutropenic mice, with mIgG_1_ having a small but significant advantage over the mIgG_3_ **(Figure 4A)**. Additionally, the mIgG_1_ appeared to promote higher expression of inflammatory cytokines IFN-γ, TNF-α, IL-12 and IL-17 in neutropenic mice, suggesting higher immune activation, while the mIgG_3_ antibody generally exhibited reductions of these markers in neutropenic mice **(Figure 4B)**.

We proceeded to compare subclass efficacy at a lethal infectious dose, which has been previously observed to cause dissemination to the liver and spleen, and death within 72 hours. At this dose, both subclasses performed equally in nondepleted mice, reducing lung, liver, and spleen burden by at least 1 log **(Figure 4C)**. However, within neutropenic mice, mIgG_1_ showed some loss of efficacy, reducing lung burden by only 0.52 log. Meanwhile mIgG_3_ continued to reduce burden in all three organs by over 1 log, though results were more variable. Cytokines levels were also more variable as expected, though global increases in inflammatory cytokines such as GM-CSF, IL-6 and IL-17 were observed compared to the non-lethal dose **(Figure S4)**. These cytokines appeared to drop in the presence of either subclass.

Overall, we observed protection of both antibodies in both immunocompetent and neutropenic mice, with a slight advantage of the mIgG_1_ subclass in mediating small localized infection. At higher doses, both subclasses demonstrated equal efficacy in healthy mice, while only mIgG_3_ subclass showed significant efficacy in neutropenic mice.

## DISCUSSION

Though the pharmaceutical industry has focused primarily on development of hIgG_1_ antibodies, there are increasing efforts to compare the efficacy of different antibody isotypes and subclasses in treating both cancer and infectious disease (31–33). Despite mIgG_2a_ and mIgG_2b_ being most similar to hIgG_1_, direct comparisons between mIgG_1_ and mIgG_3_ exclusively have provided important insights into antibody-mediated resolution of infection (12, 34, 35). Studies investigating anti-capsular antibodies against *Cryptococcus neoformans*, for example, have suggested that mIgG_3_ is poorly suited to protect against infection compared with mIgG_1_ (36, 37), whereas protection in mice from *Bacillus anthracis* spores was exclusively mediated by mIgG_3_ (8). Sub-class specific antibody interactions with *K. pneumoniae* have not been previously studied in detail. Though humoral responses to a CR-*Kp* hexasaccharide vaccine were predominantly mIgG_1_ (38), mIgG_3_ antibodies, like hIgG_2_, are the primary humoral response to T-independent antigens such as polysaccharide capsules, and the predominant monoclonal antibody subclass identified after vaccination with full length capsule (14, 39, 40). This suggests that mIgG_3_ has some evolutionary importance in protecting against encapsulated organisms.

While having identical variable regions, 17H12 mIgG_1_ and 17H12 mIgG_3_ differ in binding CR-*Kp* capsular polysaccharide, inducing serum bactericidal activity, fixing complement, promoting phagocytosis, and promoting neutrophil-mediated bacterial reduction both *in vitro* and *in vivo*. The improved binding ability of mIgG_3_ subclasses has been previously observed in antibodies against *Burkholderia* and group A *Streptococcus* capsules (35, 41) and attributed to the ability of mIgG_3_ to self-aggregate, creating opportunities for cooperative binding and increased avidity (42, 43). Our binding and agglutination data extend this knowledge to anti-capsular antibodies against CR-*Kp*, though we observed an intermediate level of binding after F(ab’)_2_ digestion of mIgG_3_ whereas others have observed mIgG_3_ F(ab’)_2_ fragments to have no better binding than mIgG_1_ fragments. (35, 41). This difference may be due to greater contribution of the mIgG_3_ hinge region or disulfide bonds to *K. pneumoniae* CPS epitopes (42), and further digestion of the antibody into Fab fragments or replacement of heavy chain domains may shed more light on these interactions (8). Nonetheless, aggregation and cooperative binding appear to be important in defending against encapsulated organisms, since in addition to mIgG_3_ and cold agglutinin IgM, the T-independent subclass hIgG_2_ has also demonstrated the ability to self-aggregate (44). Such properties however make preparations of monoclonal antibodies exceedingly difficult to purify and store.

Additionally, agglutination of the 33576 strain by our antibodies suggest clade 2 CPS functional epitopes could be conserved across geographic location. Our previous studies were restricted to *wzi154* clinical isolates collected in the greater NYC area (14, 22). Though further study of the *wzi154* and other ST258 epitopes are warranted, such evidence adds more confidence to efforts to develop cross-protective antibodies and vaccines.

Several studies have shown the resistance of CR-*Kp* strains to serum and the efficacy of antibodies in overcoming this resistance (14, 15, 22, 45). Observing serum bactericidal activity is important when comparing antibody-mediated activity against CR-*Kp*. Our results re-iterate previous findings that mIgG_3_ antibodies promote greater serum killing, as well as C3c and C5b-9 deposition, than mIgG_1_ can. Fixation of complement by mIgG_3_ has been shown to be mediated by its CH2 domain (46), and the poor ability of mIgG_1_, to fix complement has also been observed (47, 48). As previously found and also demonstrated in this study, the capsule is imperative to CR-*Kp* survival in blood (15). Thus, mIgG_3_ may be more advantageous in limiting CR-*Kp* hematogenous dissemination through complement fixation.

The relative contributions of macrophages, monocytes, and neutrophils to CR-*Kp* clearance has been disputed. While evidence using cell-specific depletions in mice strongly suggested that lung clearance of CR-*Kp* is predominantly mediated by CCR2 positive macrophages over neutrophils (26), studies examining the role of human and primate neutrophils in CR-*Kp* clearance have shown their efficacy in clearing bacteria *in vitro* (15, 29, 49). These contributions matter significantly to the field of anti-infective antibody therapy in the context of CR-*Kp*, as up to 86% of patients with CR-*Kp* bacteremia are neutropenic, and these patients have been found to fare worse prognoses than patients without neutropenia (19, 28). Our findings demonstrate that 17H12 mIgG_1_ may promote improved phagocytosis of CR-*Kp* relative to 17H12 mIgG_3_ in murine J774 cells, and similar phagocytosis in bone marrow derived macrophages. This is interesting as these antibodies are thought to act via different receptors on the macrophage surface (34, 50). Nevertheless, phagocytosis of two CR-*Kp* strains did not correlate with killing of the bacteria, as CFU and visual evidence indicated that once inside, the CR-*Kp* was able to evade killing by the macrophage, and indeed multiply within the cell. This phenomenon was also observed previously in non-CR *K. pneumoniae*, which were demonstrated to inhibit phagolysosome fusion (51). It is possible that coordination of macrophages with additional immune cells and cytokines in concert may be required for the full capability of antibody-mediated opsonophagocytosis to be realized (52–55). Additionally, alveolar macrophages or inflammatory monocytes may have improved lysosomal capabilities relative to standard bone-marrow derived macrophages (56), or macrophages may clear phagocytized bacteria self-destruction via autophagy or pyroptosis (52, 53).

Our studies in human neutrophils showed mIgG_3_ promoted better clearance of CR-*Kp* by neutrophils at lower concentrations in the presence of serum. However, while antibody-mediated killing of CR-*Kp* by neutrophils depended on sera, the production of reactive oxygen species upon stimulation with CR-*Kp* was not, as demonstrated with the production of ROS with heat-killed FBS-enriched media. Such findings suggest that ROS released by neutrophils in response to CR-*Kp* may be reactionary without being protective; further studies examining ROS responses *in vivo*, as well as studies examining other neutrophil protection mechanisms, such as lysosomal activity and neutrophil extracellular trap release, are warranted.

Our *in vivo* data provides several important findings. Firstly, as previously stated we observed CFUs in the lungs of MMC39-infected mice to be higher in the neutrophil depleted mice in both sublethal and lethal challenges, suggesting that neutrophils are indeed important in protection against pulmonary infection by this *wzi154*. This runs counter to other studies that found no change in CFU in lungs of neutropenic mice (26, 30). Our laboratory has previously discovered variability in the virulence of ST258 strains, including within *wzi154* strains (22), and additional work has determined neutrophils can clear some ST258 strains (49). Therefore, it is possible that immune responses to different CR-*Kp* strains may vary and thus be responsible for these differences.

Additionally, we observe potential differences in protection by the different subclasses. While both antibodies reduced bacterial burden in the lungs of mice, mIgG_1_ performed better and was associated with higher levels of inflammatory cytokines in neutropenic mice than the control-treated or mIgG_3_-treated mice when given a sublethal dose. We suggest that at low inocula, monocytes and macrophages may be more important for infection control, and as mIgG_1_ showed better opsonophagocytosis, it may function as the better subclass in these mice. As mentioned, bacterial control by macrophages could be augmented by other immune populations such as gamma delta T cells, and innate type III lymphocytes, all of which may produce IL-17 to potentiate *Klebsiella* immunity (30, 57, 58). Increased IL-17, IL-12 and other cytokines in the neutropenic mice may thus compensate for neutrophil responses through action by macrophages, monocytes, and other cells. IL-17, produced by resident lymphocyte populations has been identified as indispensable in protection against *K. pneumoniae* pulmonary infection (30, 57–60). However, these neutrophil-independent responses may be insufficient at higher inoculums, as evidenced by the reduction of mIgG_1_ efficacy in the neutropenic mice challenged with lethal infection. In contrast, drops in inflammatory cytokines in mIgG_3_ treated neutropenic mice may indicate other components, such as complement, may be sufficient to control infection and require fewer compensatory distress cues for local control. Complement has been shown to be important in lung clearance on other pathogens early in infection (61, 62). Furthermore, retention of efficacy by mIgG_3_ in the high inoculum, neutropenic scenario may suggest a large role of complement in defending against more disseminated infection, when local control of infection by macrophages and other resident populations may be insufficient to compensate for the role of neutrophils. Studying subclass using complement-depletion models may provide additional insights into the relative contribution of complement in antibody-mediated protection.

Our study has several limitations. Firstly large heterogeneity in virulence exists between *K. pneumoniae* isolates, even within CR-*Kp* subsets (22), and our contrasting findings regarding neutrophil protection highlight the need to further study heterogeneity of the pathogen-immune response in numerous CR-*Kp* isolates (26). Furthermore, the functions of individual monoclonal antibodies, even those with the same subclass and similar target, can also be heterogeneous. Some anti-capsular *Streptococcus pneumoniae* antibodies are better able to promote opsonophagocytosis, while others may directly interference with bacterial signaling and growth (63, 64). Therefore, future studies of CR-*Kp* anti-capsular antibodies should investigate several to better sample their potential. Finally, future studies of anti-CR-*Kp* antibodies must innovate the field of *in vivo* models used to study CR-*Kp* infection, as convenient and effective models that reproduce the chronic, persistent infection caused by CR-*Kp* in humans have been difficult to develop (29).

In conclusion, we find that the subclass differences of an anti-capsular antibody can affect various facets of immune function against carbapenem-resistant *Klebsiella pneumoniae*, but can exhibit similar efficacy *in vivo*. This information will promote future monoclonal antibody work on CR-*Kp* to provide effective therapies, and supports the potential of antibody 17H12 as a candidate to further study. Furthermore, we observe a role for neutrophils in antibody-mediated protection *in vivo*, encouraging further efforts to investigate the pathogen-host interactions of CR-*Kp*.

## METHODS

### Ethics Statement

Animal study protocols were approved by the Animal Committee (IACUC) at Stony Brook University (Protocol 628253) in accordance with the Guide for the Care and Use of Laboratory Animals, the Animal Welfare Act, Public Health Service Policy on Human Care and Use of Laboratory Animals, and all other local, state and federal regulations. Healthy serum and neutrophil donors gave written informed consent for blood donation under IRB protocol 718744 at SBU.

### Bacteria and Growth Conditions

*Klebsiella pneumoniae* (MMC)5, (MMC)34, and (MMC)39 are *wzi154* clinical isolates, and (MMC)36 and (MMC)38 are *wzi29* and *wzi50* isolates, respectively, from Montefiore Medical Center and described previously (22). SBU32 and SBU34 are clinical isolates from Stony Brook University Hospital previously used to study 17H12 mIgG_3_ (14). 33576 and its capsule-deficient mutant (33576Δwzy) were graciously provided by Dr. Barry Kreiswirth (15). For all experiments (unless otherwise noted), strains were grown at 37°C shaking to mid-exponential phase in Miller LB broth from a 1:100 dilution of an overnight culture. Overnight cultures were derived from single colonies picked from a Miller LB plate no older than thirty days. Unless stated otherwise, cultures were washed with PBS twice before use.

### Generation of pProbe-KtBl and Transformation of CR-*Kp* MMC39

We previously utilized a pPROBE-Kt GFP plasmid with a pVS1 backbone and with GFP under control of an inserted *nptII* Kanamycin promoter (27, 65). We used this plasmid, and under contract with Genewiz inserted the sequence of the *sh ble* gene, which conveys resistance to bleomycin, into a ClaI cut site positioned between the *ori* and the existing *kan^R^* cassette. Insertion of the gene was confirmed both by sequencing and by digest with BssHII, which both the original pPROBE-Kt and the *sh ble* gene possessed. The plasmid was transformed into MMC39 by electroporation and plated onto Lennox (low salt) LB, pH adjusted to 8.0, containing 50μg/mL Bleomycin (Zeocin™, ThermoFisher). Transformation was confirmed by growing positive colonies on 50μg/mL bleomycin, 50μg/mL kanamycin, and observing GFP fluorescence of selected colonies under a microscope. To test for plasmid stability, we tested replicate plating of 100 unselected colonies onto selective plates, all of which grew. Additionally, nearly all bacteria screened after ten serial exponential cultures of MMC39 in the absence of antibiotics were shown to be GFP+ by microscopy. For all later experiments MMC39-GFP isolates were streaked onto Miller LB agar supplemented with 50ug/mL of kanamycin and grown in kanamycin-supplemented Miller LB broth.

### Subclass Switch Production and Sequencing

The mIgG_1_ switch variant of the 17H12 mIgG_3_ murine hybridoma was generated as previously described (12). Briefly, the mIgG_3_ parent hybridoma line was treated with endotoxin and IL-4, and spontaneous switch variants were identified through ELISPOT. Sib selection was initially utilized, followed by two rounds of fluorescence-activated cell sorting (FACS) of cells stained with FITC-Conjugated rat anti-mouse IgG_1_ (Southern Biotech 1144-02), and rounds of soft-agar cloning in SeaPlaque agarose. Vials of frozen hybridoma clones were sent to GenScript for variable region exon sequencing and IMTG analysis.

### Antibody Purification

Antibodies were produced weekly over six months from respective hybridomas grown in CELLine (Wheaton) flasks fed with High-Glucose DMEM + 10% NCTC media and 1x Penicillin-Streptomycin and 1x Non-Essential Amino Acids, supplemented with either 10% or 5% FBS in the inner and outer chambers, respectively. These antibodies were purified using Pierce Protein G Affinity Chromatography as per manufacturer’s instructions. Eluted antibody was neutralized in Tris-HCl, pH 8.0, and NaCl to final concentrations of 100mM and 300mM, respectively, then concentrated by centrifugal filtration (AMICON 30K), filter-sterilized, snap frozen in liquid nitrogen, and stored at −80° C until use. Concentration was determined by absorbance at 280nM (Extinction coefficient = 1.4), which correlated with Bradford assay results.

### F(ab’)_2_ generation

F(ab’)_2_ fragments of 17H12 mIgG_3_ and mIgG_1_ were generated and purified using the Pierce F(ab’)2 Preparation Kit and Mouse IgG_1_ Fab/F(ab’)2 Preparation Kits following manufacturer’s instructions, except for digestion temperature and duration (10 min at ambient temperature for mIgG_3_, 36 hours at 37°C for mIgG_1_). Coomassie staining of SDS-PAGE in nonreducing conditions was utilized to ensure mAb/F(ab’)_2_ purity after all purifications/digestions.

### Binding Affinity

The EC_50_ of the mAbs was calculated using ELISA as described previously (27). Briefly, polystyrene plates (Corning 3690) were coated with 0.5mg/mL of ST258 clade 2 CPS (MMC34) in PBS, then blocked with 1% PBS-BSA. The mAbs or F(ab’)_2_ fragments were serially diluted starting at 1 μM (assuming a MW of 150kD for full IgG and 110kD for F(ab’)_2_) and proceeding twofold. Antibody was detected using an HRP-conjugated Goat anti-Mouse IgG kappa secondary antibody (Southern Biotech, PA1-86015, 1:1000) and developed with 1-Step Turbo TMB ELISA Substrate (ThermoFisher) according to manufacturer’s instructions. Between steps wells were washed four times with PBS 0.1% Tween-20. Experiments were repeated on two different days with two different antibody purification batches and digests to ensure reproducibility. Control antibodies were run in parallel as negative controls (Southern Biotech 0102-01, 0105-01).

### Agglutination

Agglutination of bacteria by mAbs was detected by flow cytometry and confirmed by microscopy similar to previous studies (25, 66). Washed cultures were diluted to approximately 1 x 10^8^ CFU/mL, and 40 microliters were added to 160uL of PBS with the appropriate antibody concentration and 0.5% BSA to give a final concentration of 2 x 10^7^ CFU/mL. The samples were mixed gently in round-bottom flow cytometry tubes and incubated at 37°C in a shake incubator for 1 hour, and later fixed by adding 200uL of 2% PFA and incubating for 10 min at RT. Fixed samples were analyzed via FACS Calibur using Forward and Side Scatter to determine clump sizes, and also visualized on glass slides under phase contrast microscopy (EVOS FL Auto, 40x and 100x Objective, ThermoFisher). Voltages for flow cytometry were fixed throughout, but gating of bacteria was adjusted based on individual strains incubated in PBS alone to account for size differences between strains. 50,000 events within the gate were counted per sample, representing ~75% of all recorded events. The percentage of positive agglutination events was calculated by measuring the percentage of gated cells whose forward scatter exceeded a value representing the largest 1% of events for bacteria treated with PBS alone. Control antibody 14G8 utilized was a mIgG_1_ against *Staphylococcus aureus* Enterotoxin B (SEB) (67).

### Serum Resistance Assays

Serum resistance/killing assays were modified from a previously described assay (22). Briefly, 250mcL of a 1x 10^5^ CFU/mL solution was added to 750mcL of PBS containing 20% fresh or heat-inactivated (HI) human serum from a healthy donor. HI serum was generated from donor serum by 30 min incubation in a 57°C water bath. Tubes were incubated at 37°C rotating end-over-end. At 0 min, 60 min, and 120 min after mixing, 100μL was sampled from each tube, diluted, and plated onto LB agar for CFU quantitation. Percent survival was measured as a fraction of the CFU count at 0 min.

### Complement Deposition Assays

Flow cytometry was used to detect complement deposition of C3c and C5b-9, as previously described (14). Briefly, bacteria were diluted to 1 x 10^8^ CFU/mL in 1mL PBS-BSA 1% or 20% fresh human serum (in PBS-BSA). PBS or 10μg/mL of antibody was then added, and bacteria were incubated either 20 min or 40 min at ambient temperature for C3c and C5b-9 deposition, respectively. Bacteria were washed, resuspended in PBS-BSA, and incubated with either FITC-conjugated Sheep Anti-Human C3 (BioRad AHP031F) at 1:500 or AF488-conjugated Mouse Anti-C5b-9 (ae11) (Novus Biologicals 5120AF488) at 1:150, or without antibody for 20 min at 4°C. After incubation bacteria were washed and analyzed for fluorescence by FACS Calibur. Integrated geometric mean fluorescence was measured as the product of the percent of gated events that passed a fluorescence threshold and the mean fluorescence of those events that passed the threshold.

### Macrophage Phagocytosis Assays

BMDM were differentiated from frozen bone marrow from 6 week-old c57BL/6 mice (Taconic) as previously described (68), except using pure M-CSF (10ng/mL) rather than L929 media as the M-CSF source for feedings on Day 1 and 4. Differentiation of cells was confirmed by flow cytometry on cells stained with FITC-conjugated anti-F4/80 and BV510-conjugated CD11b (purity >98% double-positive) Macrophage phagocytosis as measured by CFU was performed similarly to previous protocols (22, 27), Briefly, 1×10^5^ BMDM or J774A.1 cells were incubated overnight in wells of cell-culture treated 96 well plates in either RPMI with HEPES and L-Glutamine + 10% FBS +1x Non-Essential Amino Acids; or High-Glucose DMEM + 10% FBS +10% NCTC-109 +1x Non-Essential Amino Acids; respectively, at 37°C in 5% CO_2_. The following day 1×10^7^/mL bacteria were opsonized for 20 min in respective cell-culture media containing either 40μg/mL of mIgG_1_, mIgG_3_, or control mIgG_1_, and 100mcL of this (MOI 10) was added to each well of the washed macrophage plates. After 30 min incubation at 37°C + 5% CO_2_ cells were washed thrice and exposed to media with 100μg/mL of Polymyxin B for 20 min. Cells were washed again, and Time 0 wells were immediately lysed twice with water and plated, while later time points remained in culture media until needed. All conditions were performed in triplicate wells. The number of CFU calculated from LB plates was divided by the number of estimated cells plated to give the phagocytic index. Microscopy was performed using the MMC39-GFP strain and EVOS FL Auto, 40x Objective, ThermoFisher using phase contrast and a GFP light cube.

### Neutrophil Killing Assays

Neutrophil assays were adapted from a previous protocol (15). Briefly, 5 x 10^5^ bacteria opsonized 30min at RT in RPMI containing appropriate antibody concentrations were added to 5 x 10^5^ human neutrophils in RPMI containing a final concentration of 5% autologous fresh or HI serum. At 0 min, 15 min, 30 min, and 60 min, 100mcL were sampled from reaction tubes, lysed in cold PBS + 0.1% Triton X-100, diluted in PBS, and plated on LB agar for CFU quantitation. Percent survival was measured as a fraction of the CFU count at 0 min. Tubes containing sera but not neutrophils were run in parallel. Neutrophils were purified from whole blood by MACSxpress^®^ Whole Blood Neutrophil Isolation Kit (Miltenyi), treated once for 5 min with red blood cell lysis buffer, and resuspended in RPMI on ice until use. Flow cytometry performed on neutrophils purified from two experiments both showed purity >99%.

### Pulmonary Infection Experiments

We used c57BL/6 mice (Taconic), aged 7-9 weeks for all mouse experiments, and pulmonary infection was performed as previously (14, 69). At 48 hours and 4 hours prior to the procedure, mice were injected intraperitoneally with 225ug of Rat anti-mouse Ly6G (1A8) or a control Rat anti-mouse IgG_2a_ (2A3) (BioXcell). Neutrophil depletion was confirmed previously using flow cytometry of lung homogenates (Ly6G^+^, Ly6C^-^, CD11b^-^). Inoculums were prepared by resuspending MMC39 in a Tris-Glycine buffer containing with 5mg/mL of an ovalbumin control mIgG_1_ (Crown Biosciences), 17H12 mIgG_1_, or 17H12 mIgG_3_, to a final concentration of 6×10^6^ or 3×10^7^ CFU/mL. After 1-hour opsonization, 50mcL of the inoculum was instilled into the surgically-exposed trachea of a mouse under ketamine/xylazine using a bent 27G needle. After 20 hours, mice were euthanized and lungs, liver, and spleen, were collected and processed in NP-40 or PBS and diluted to enumerate CFUs. Supernatants of lung homogenates used for cytokine analysis were stored at −80 with 1x Pierce Proteinase Inhibitor until testing using Bio-Plex Pro Mouse Cytokine Th17 Panel A with additional GM-CSF and IL-12p70 Singleplex sets on a Bio-Plex200 Platform (BioRad). Cytokine levels were normalized against total protein measured by Bradford assay.

---

**<u>Figure S1.</u>** In-depth comparison of agglutination of 17H12 mIgG_1_ and mIgG_3_. (A) Comparison of bacterial population agglutination sizes of MMC39-GFP incubated with various concentrations of 17H12 mIgG_1_ (left, red), or 17H12 mIgG_3_ (right, blue). The geometric mean forward scatter of the entire GFP-positive population is shown for each concentration. (B-E) Representative agglutination plots of MMC5 (B,D) and 33576 (C,E). B and C depict gating scheme and representative microscopy images at 40x magnification, while C and D depict representative data from one of three independent experiments for each bacterial strain. Forward scatter means of both the entire gated population and the percent positive marker (back bar in histogram) are displayed. F. Bar graph of the Geometric Mean Forward Scatter of the entire gated population of bacteria agglutinated with control antibody, either 17H12 antibody, or PBS alone. Two-Way ANOVA with Tukey’s posthoc test found significant differences between PBS and either 17H12 Ab (p < 0.001), as well as between Control and either 17H12Ab (p < 0.001) for MMC5, MMC34, MMC39, SBU32, SBU34, and 33576, and no difference between any treatments groups for 33576Δwzy, MMC36, or MMC38.

**<u>Figure S2.</u>** Comparison of binding of 17H12 mIgG_1_ and 17H12 mIgG_1_ F(ab’)_2_ (A) Binding curves of 17H12 mIgG_1_ and its F(ab’)_2_ measured by indirect ELISA on plates coated with *wzi154* (MMC34) CPS. EC_50_ values are displayed with legend. The plot is representative of one experiment performed in triplicate. As the secondary antibody used in this ELISA was anti-IgG, as opposed to anti-Kappa (light chain), OD maxima differed in the detection of the two antibodies. Therefore, OD maxima were normalized to 100% for this experiment alone.

**<u>Figure S3.</u>** Serum Bactericidal Effect and Complement Deposition Controls (A, B) Growth curves of 33576 in 40% NHS supplemented (A), or 33576Δwzy in 20% NHS (B) with 40μg/mL of indicated mAb. 100% represents no increase in CFU from baseline. Error bars indicate SEM of the number of experiments noted in the graph. (C, D) Fixation of complement components onto MMC39 (top), and 33576 Δwzy (bottom), as measured by flow cytometry, with the addition of 1 experiment using HI-Serum per complement component in MMC39. Bars indicate mean and SEM of three independent experiments unless otherwise noted. MMC39 Data in C and D for Serum and Non-Serum conditions are reproduced from data in Figure 2 C & D to show contrast with the HI-Serum.

**<u>Figure S4.</u>** Reactive Oxygen Species production by human neutrophils exposed to pre-opsonized MMC5 (A,B), MMC32 (C,D), and MMC34 (E,F), as measured by luminol luminescence, in the presence of NHS (A,C,E) or FBS (B,D,F). The left time lapse graphs are representative of at least three independent experiments. Right bar graphs show aggregate data of the Area Under the Curve (A.U.C.) and the maximum rate of change (Max Δ), relative to PBS, for all experiments. Differences in A.U.C. and Max Δ between control Ab, mIgG_1_ and mIgG_3_ were assessed for significance by a Kruskal-Wallis test, with results of Dunn’s test for multiple comparisons displayed in-graph. For all in-graph statistics, p values displayed in black are comparisons to the control IgG, whereas p values in red compare mIgG_1_ with mIgG_3_. p values are replaced with ns if >0.1 (not significant); * if < 0.05; and ** if <0.01

**<u>Figure S5.</u>**Lung cytokine levels of mice that received lethal pulmonary challenge. Cytokine levels displayed are normalized to total lung protein, For all studies, overall differences in CFU and cytokines between treatment groups and between neutrophil status were assessed for significance by Two Way-ANOVA. Individual comparisons made between treatment groups of mice of the same neutrophil status (* symbols above), were tested using Tukey’s post-hoc test with p values displayed in-graph. p values are replaced with ns if >0.1; * if < 0.05; ** if <0.01; and *** if p<0.001. p values below plots compare CFU or cytokine levels of wild type and neutropenic mice within the same treatment group.

## Acknowledgements

This grant was funded using NIAID F30 AI140611 and US Veterans Affairs Merit Review Award I01 BX003741. BCF is an attending at the U.S. Department of Veterans Affairs - Northport VA Medical Center, Northport, NY. The contents of this review do not represent the views of VA or the United States Government. We would like to thank Liang Chen and Barry Kreiswirth for gifting us CR-*Kp* strain 33576 and its acapsular mutant. We would also thank Bruna Seco and Peter Seeberger for their initial contributions to the project. Additionally, we would like to thank Anne Savitt for her assistance in editing the manuscript. Authors report no financial conflicts of interest.

